# Maternal Obesity and Western-style Diet Impair Fetal and Juvenile Offspring Skeletal Muscle Insulin-Stimulated Glucose Transport in Nonhuman Primates

**DOI:** 10.1101/864082

**Authors:** William Campodonico-Burnett, Byron Hetrick, Stephanie R. Wesolowski, Simon Schenk, Diana L. Takahashi, Tyler A. Dean, Elinor L. Sullivan, Paul Kievit, Maureen Gannon, Kjersti Aagaard, Jacob E. Friedman, Carrie E. McCurdy

**Affiliations:** Department of Human Physiology, University of Oregon, Eugene, OR, USA; Department of Biochemistry, University of Colorado Boulder, Boulder, CO, USA; Department of Pediatrics, Perinatal Research Center, University of Colorado, Aurora, CO, USA; Department of Orthopaedic Surgery, University of California San Diego, La Jolla, CA, USA; Biomedical Sciences Graduate Program, University of California San Diego, La Jolla, CA, USA; Division of Cardiometabolic Health, Oregon Health Science University, Oregon National Primate Research Center, Beaverton, OR, USA; Division of Neuroscience, Oregon Health Science University, Oregon National Primate Research Center, Beaverton, OR, USA; Department of Medicine, Division of Diabetes, Endocrinology, and Metabolism, Vanderbilt University Medical Center, Nashville, TN USA; Department of Obstetrics and Gynecology, Division of Maternal-Fetal Medicine, Baylor College of Medicine, Houston, TX, US; Harold Hamm Diabetes Center, University of Oklahoma Health Sciences Center, Oklahoma City, OK, 73104 USA

**Keywords:** developmental programming, insulin signaling, glucose uptake

## Abstract

Infants born to mothers with obesity have a greater risk for childhood obesity and metabolic diseases; however, the underlying biological mechanisms remain poorly understood. We used a nonhuman primate model to investigate whether maternal obesity combined with a western-style diet (WSD) impairs offspring muscle insulin action. Briefly, adult females were fed a control (CON) or WSD prior to and during pregnancy and lactation. Offspring were weaned to a CON or WSD. Muscle glucose uptake and insulin signaling were measured *ex vivo* in fetal and juvenile offspring. *In vivo* signaling was evaluated before and after an intravenous insulin bolus just prior to weaning. We find that fetal muscle exposed to maternal WSD had reduced insulin-stimulated glucose uptake and impaired insulin signaling. In juvenile offspring, insulin-stimulated glucose uptake was similarly reduced by both maternal and post-weaning WSD. Analysis of insulin signaling activation revealed distinct changes between fetal and post-weaning WSD exposure. We conclude that maternal WSD leads to a persistent decrease in insulin-stimulated glucose uptake in juvenile offspring even in the absence of increased offspring adiposity or markers of systemic insulin resistance. Switching offspring to a healthy diet did not ameliorate the effects of maternal WSD suggesting earlier interventions may be necessary.

## Introduction

Obesity and associated metabolic disorders, including type 2 diabetes and cardiovascular disease, are major health issues worldwide, reaching epidemic proportions in western populations. In the United States, obesity continues to rise steadily with recent reports that 39.8% of adults are obese (1). Reflecting this trend is the alarming rise in childhood obesity. While lifestyle factors clearly contribute to obesity, a growing body of data finds that maternal obesity or a western style diet (WSD) rich in saturated fat during pregnancy has adverse long-term metabolic effects on the offspring (2). Population based studies demonstrate that obesity during pregnancy is associated with an almost 4-fold greater risk of childhood obesity (3) and a 2-fold higher risk of developing metabolic syndrome in adolescents born to mothers with obesity (4–6). Genome wide association studies find that only a fraction of intergenerational predisposition for obesity can be explained by genetic polymorphisms (7), suggesting that environmental exposures during development can program offspring metabolism later in life.

The developmental origin of health and disease (DOHaD) hypothesis postulates that exposure to environmental challenges during critical windows of development results in fetal adaptations that become maladaptive when exposed to subsequent metabolic and environmental stressors (8). Consistent with DOHaD, mesenchymal stem cells from infants exposed to maternal obesity have a greater propensity towards adipocyte versus myocyte differentiation, reduced lipid oxidative capacity when challenged with lipid and hypermethylation of genes regulating fatty acid oxidation, demonstrating functional consequences downstream of environmental stressors during development (9,10).

Skeletal muscle insulin resistance is a primary defect in the etiology of type 2 diabetes (11,12). Thus, it is important to understand the impact of maternal diet on skeletal muscle insulin response, especially in young offspring. Multiple studies find that developmental exposure to maternal obesity or a maternal diet high in saturated fats during gestation predisposes offspring to obesity and insulin resistance (13); however, the mechanism by which early life exposure to maternal obesity and WSD may alter skeletal muscle insulin-stimulated glucose transport and signal transduction is relatively untested.

We used an established non-human primate model to investigate the impact of maternal obesity and a WSD on programming of offspring skeletal muscle insulin action (14–17). We hypothesized that exposure to maternal obesity concomitant with WSD feeding during fetal development and lactation would impair insulin-stimulated glucose uptake in offspring skeletal muscle independent of offspring adiposity and the effect would be exacerbated by a post-weaning WSD. To test our hypothesis, we measured insulin-stimulated glucose uptake and canonical insulin signaling in isolated muscle strips from fetal and juvenile offspring from lean control (CON) fed dams or obese WSD-fed dams, with cross-over post-weaning diets among the juvenile offspring (Figure 1).

**Figure 1.**
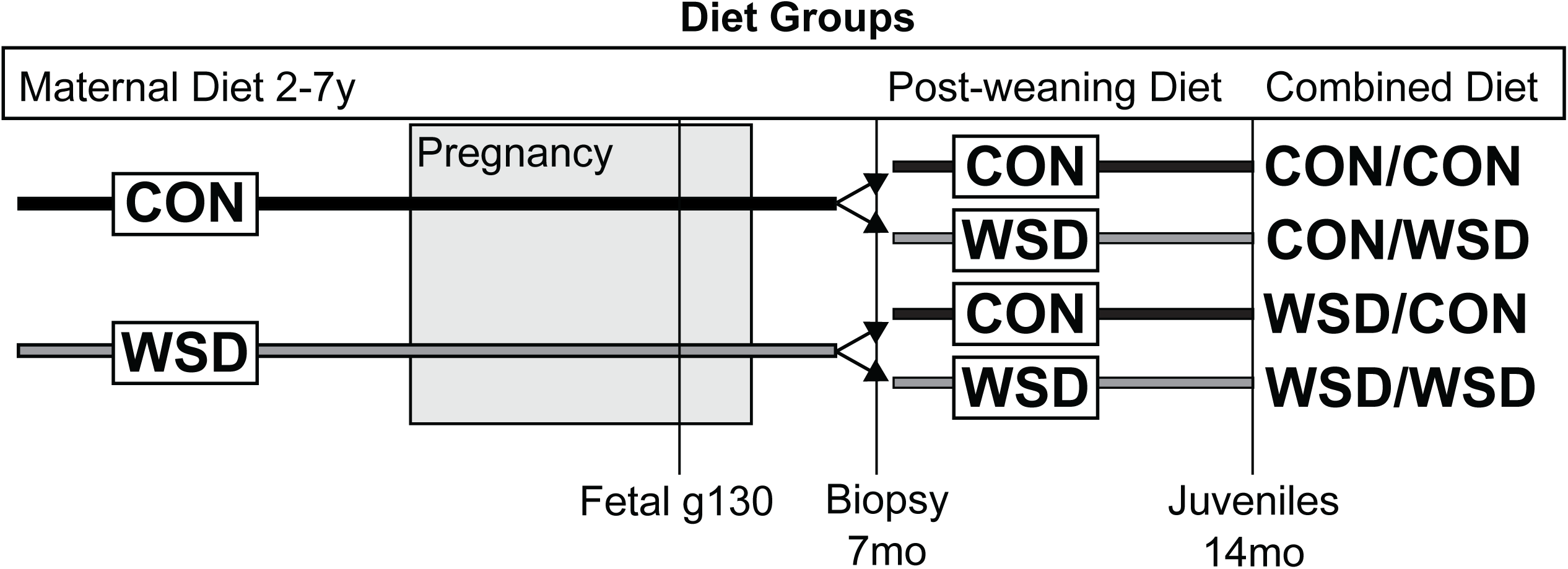
Schematic of Experimental Design. Female Japanese Macaques were placed on CON or WSD for 2-7 years prior to pregnancy and maintained on the same diet through pregnancy and weaning of offspring. Post-weaning, offspring were either maintained on the same diet or switched to the alternate diet. Fetal samples were assayed at gestational day 130, juveniles were assayed at 14 months of age, and skeletal muscle biopsies were assayed from animals at 7 months of age.

## Research Design and Methods

### Experimental Animal Model

All animal procedures were approved by and conducted in accordance with the Institutional Animal Care and Use Committee of the Oregon National Primate Research Center (ONPRC) and Oregon Health and Science University. The ONPRC abides by the Animal Welfare Act and Regulations enforced by the USDA and the Public Health Service Policy on Humane Care and Use of Laboratory Animals in accordance with the Guide for the Care and Use of Laboratory Animals published by the NIH.

Adult Japanese macaques were group-housed in indoor-outdoor enclosures and fed ad libitum CON diet containing 15% calories from fat primarily from soybeans and corn (Monkey Diet no. 5052; Purina Mills) or WSD with 36.6% calories from fat primarily from animal fat, egg, and corn oil (TAD Primate Diet no. 5LOP; Purina Mills) for 2-7 years. The carbohydrate sources differed between the two diets, with sugars (primarily sucrose and fructose) comprising 19% of the WSD but only 3% of the CON. All animals fed the WSD were also given calorically dense treats once per day. Females were allowed to breed seasonally, and gestational age was determined by ultrasound (18,19). Maternal body fat percent was determined by DEXA in non-pregnant females 3 months prior to the start of the breeding season (17). Intravenous glucose tolerance tests (IVGTT) were performed on adult females in the non-pregnant state (3 months prior to the start of the breeding season) and during the 3rd trimester of pregnancy as previously described (14,20). A detailed characterization of maternal metabolic profiles and maternal-fetal plasma measures, as well as changes in the placenta and fetal pancreas and liver have been described elsewhere (20,21). For fetal sample collection, dams were fasted overnight, anesthetized and fetuses were delivered by Cesarean section at gestational day 130 (g130) of 165.

A subset of juvenile offspring was delivered naturally and remained with the maternal group and on the maternal diet until weaning at 6-8 months of age. At weaning, juvenile offspring from both maternal diet groups were combined into new social groups and housed in enriched, indoor-outdoor units with 6–10 similarly aged juveniles and 1–2 unrelated adult females per group. Juvenile offspring were fed CON or WSD post-weaning with nutritional supplements as described above until necropsied at 14 months of age. One week prior to necropsy, fasting juvenile offspring underwent sedated IVGTT using 0.6 g/kg glucose. Baseline and timed samples of glucose and insulin were taken. Glucose and insulin AUC were calculated in Graphpad Prism v. 8.2 using fasted basal blood glucose as baseline.

### Offspring Insulin-stimulated Biopsy Collection

Prior to weaning, at approximately 7 months of age, biopsy samples were collected from the soleus of fasted animals under anesthesia. Briefly, animals were fasted for 5 hours in the morning, sedated with telazol (4mg/kg), intubated and an intravenous catheter placed in each arm. Using an open biopsy technique, a baseline sample (∼30mg) was dissected from the soleus. Insulin (Humulin R, Eli Lily) was injected intravenously at 0.05 U/kg and a second muscle biopsy was taken 10 minutes later. Biopsy samples were snap-frozen in liquid nitrogen and stored at −80°C.

### Fetal and Juvenile Tissue Collection

After euthanasia, fetal g130 and 14-month-old offspring skeletal muscles including gastrocnemius (gastroc), soleus, vastus lateralis and rectus femoris were removed, flash frozen in liquid nitrogen, and stored at −80°C. The contralateral soleus and rectus femoris in the fetus and the soleus and gastrocnemius in the juvenile were clamped with paired hemostats at a fixed distance prior to dissection and clamped muscle was placed in ice-cold oxygenated (95% O2, 5% CO2) flasks of Krebs-Henseleit buffer (KHB). Clamped muscle was transported from necropsy to the adjacent laboratory for *ex vivo* analysis of muscle glucose uptake.

### *Ex vivo* 2-deoxyglucose (2DG) Uptake

Insulin-stimulated 2DG uptake was measured in isolated skeletal muscle strips from fetal and juvenile offspring at necropsy using a modified protocol from (22,23). Briefly, muscles were clamped in situ with paired hemostats that were mounted at a fixed distance of 2 or 3 inches. The clamped region was cut away from the intact muscle and the muscle held within the hemostat was immediately placed in ice-cold oxygenated (95% O2, 5% CO2) flasks of KHB. Fascia was scraped from the outer surface of the muscle tissue. Thin muscle strips (1 or 0.5 inches in length) were rapidly isolated from the clamped muscle, fastened with 2-prong clips and excised. A total of 3-9 muscle strips were isolated. Clipped muscle strips were pre-incubated at 35°C for 30 min in vials with oxygenated KHB containing 0.1% BSA, 2 mM Na-pyruvate, and 6 mM mannitol and either no insulin or insulin (Humulin-R, Lilly, Indianapolis, IN) at 0.3nM (sub-maximal), or 12nM (maximally stimulated). After 30 min, muscles were transferred to a second vial and incubated at 35°C for 20 min in KHB plus 0.1% BSA, 9 mM [^14^C]-mannitol (0.025 mCi/mmol; Perkin Elmer, Boston, MA), and 1 mM [^3^H]-2DG (2 mCi/mmol; Perkin Elmer, Boston, MA) with the same insulin concentration. After 20 min, muscles were trimmed from clips, blotted on ice-cold filter paper, and snap-frozen in liquid nitrogen. Muscles were weighed then homogenized in 0.5 mL cell lysis buffer (20mM Tris-HCl pH 7.4, 150mM NaCl, 20mM NaF, 2mM EDTA pH 8.0, 2.5mM Na4P2O7, 20mM ß-glycerophosphate, 1% NP-40, 10% glycerol with protease and phosphatase inhibitor cocktails using glass-on-glass tissue grinders (Kontes). Samples were solubilized (1h, 4°C with end over end rotation), centrifuged (12,000 × g), and the supernatant was collected. 0.1 mL of supernatant was quantified by scintillation counting. The rate of 2DG uptake was calculated as previously described (24,25).

### Protein Analysis

Protein concentration of muscle homogenates was determined by Pierce BCA Assay according to manufacturer’s instructions. Abundance and activation of insulin signaling proteins were measured by Simple Western (WES, Protein Simple, San Jose, CA). Plates were loaded according to manufacturer’s instructions with samples at a final concentration of 0.2 mg/mL total protein. Primary antibody dilutions used are reported in Supplementary Table 1. Proteins were separate using the 12-230 kDa capillary separation module (Protein Simple, #SM-W003) data was quantified using Compass Software v4.0.

### Statistical Analysis

Data was analyzed as indicated in figure legends using Graphpad Prism v8.2.

## Results

### Dam phenotype

The characteristics of a larger cohort of dams on WSD has previously been published elsewhere (16). Compared to dams fed a CON diet, WSD fed dams had increased body weight, body fat percent, and elevated fasting insulin both pre-pregnancy, and during the 3^rd^ trimester (Table 1). Specifically, WSD resulted in obesity defined as body fat percentage above 30%. Insulin area under the curve (IAUC) was significantly elevated pre-pregnancy and during the 3^rd^ trimester IVGTT, but the glucose AUC was comparable. Together, these findings indicate insulin resistance in the WSD fed maternal group.

**Table 1:** Maternal Phenotype.

| Parameter | CON<br>Average | SEM | n | WSD<br>Average | SEM | n | P |
| --- | --- | --- | --- | --- | --- | --- | --- |
| Body fat % | 13.9 | 1.5 | 12 | 36.2 | 1.8 | 17 | **<0.0001** |
| Weight (pre-pregnancy) (kg) | 8.8 | 0.5 | 12 | 13.8 | 0.6 | 17 | **<0.0001** |
| Weight (3<sup>rd</sup> trimester) (kg) | 9.8 | 0.6 | 11 | 13.8 | 0.6 | 13 | **<0.0001** |
| Pre-pregnancy Fasting Insulin (mU/L) | 13.6 | 2.8 | 12 | 33.7 | 7.0 | 17 | **0.03** |
| 3<sup>rd</sup> Trimester Insulin (mU/L) | 14.1 | 3.4 | 11 | 37.1 | 8.9 | 12 | **0.03** |
| Pre-pregnancy Fasting Glucose (mg/dL) | 57.1 | 3.3 | 12 | 54.4 | 1.5 | 17 | 0.4 |
| 3<sup>rd</sup> Trimester Glucose (mg/dL) | 46.6 | 2.1 | 11 | 44.8 | 2.3 | 10 | 0.6 |
| Pre-pregnancy IAUC | 1890 | 388 | 12 | 6365 | 692 | 17 | **<0.0001** |
| 3<sup>rd</sup> Trimester IAUC | 2419 | 364 | 10 | 7962 | 954 | 11 | **<0.0001** |
| Pre-pregnancy Glucose AUC | 6316 | 469 | 12 | 6879 | 400 | 17 | 0.3 |
| 3<sup>rd</sup> Trimester Glucose AUC | 5193 | 376 | 11 | 4851 | 310 | 11 | 0.9 |

### Maternal obesity and WSD reduces insulin-stimulated glucose uptake in fetal skeletal muscle

To test the effect of maternal WSD on insulin-stimulated glucose uptake, [H^3^] labeled 2- deoxyglucose (2DG) uptake into skeletal muscle was measured in the presence of low (0.3 nM) insulin, high (12 nM) insulin, or in the absence of insulin (basal). In rectus femoris, there was no significant difference in basal glucose uptake between maternal CON and WSD fetuses (Figure 2A). Insulin-stimulated 2DG uptake, expressed as a fold increase over basal, was significantly reduced in rectus femoris and soleus from WSD compared to CON at both insulin concentrations (Figure 2B, D). In the soleus, there was a significant increase in basal 2DG uptake with maternal WSD (Figure 2C).

**Figure 2:**
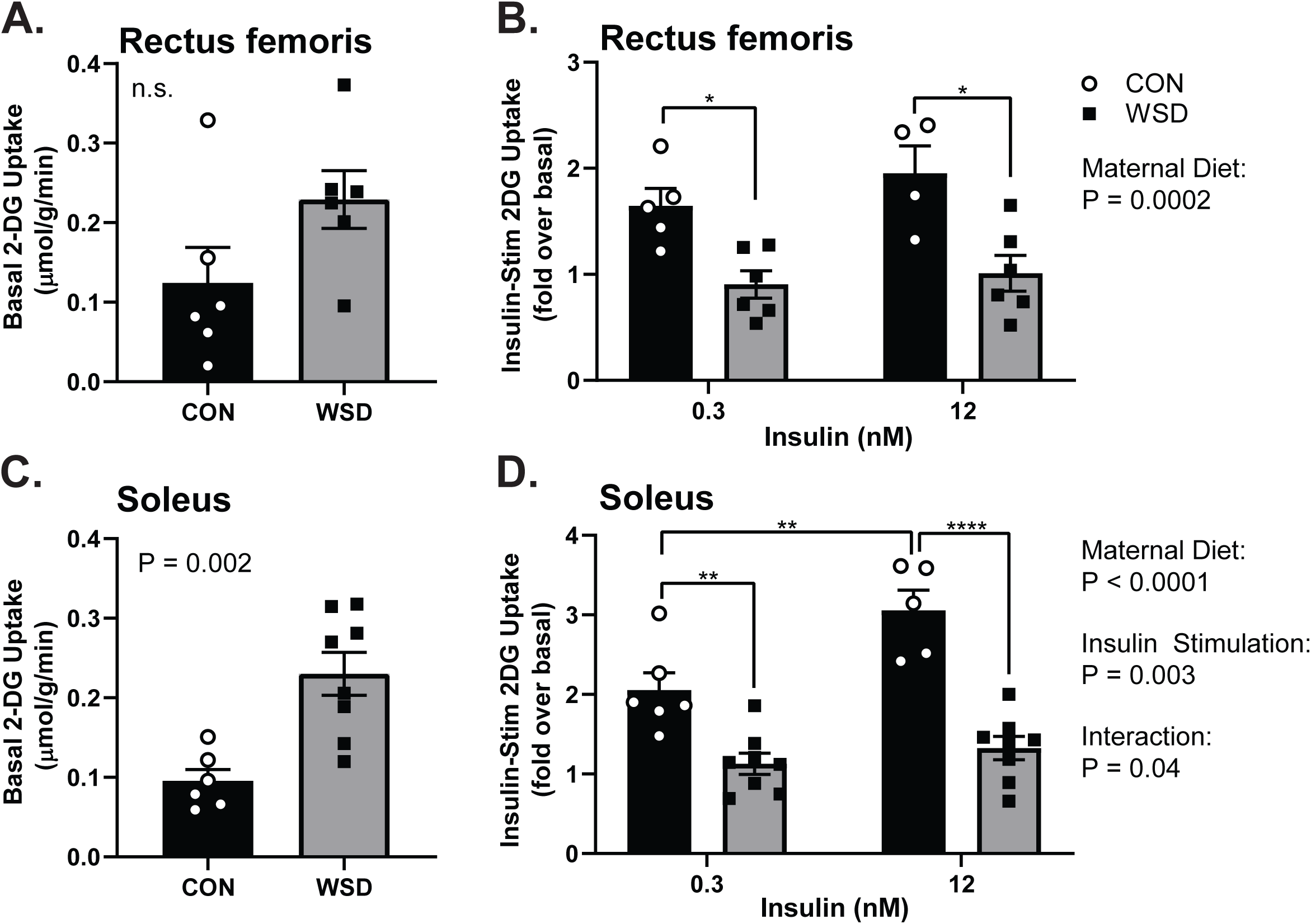
Fetal Skeletal Muscle *Ex Vivo* Glucose Uptake. Muscle strips from male and female fetal offspring at gestational day 130 from dams on CON or WSD diet were assayed for *ex vivo* glucose uptake. 2-deoxyglucose (2DG) uptake was measured in basal (A, C) and insulin-stimulated (B, D) rectus femoris (A, B) and soleus (C, D) muscles at 0.3 nM and 12 nM insulin doses. Data is expressed as mean ± SEM, with individual data points shown. Basal glucose uptake was analyzed by unpaired T test. Insulin-stimulated glucose uptake was analyzed by two-way ANOVA with Tukey multiple comparisons test. Brackets indicate group comparisons (* P≤0.05, ** P≤0.01, *** P≤0.001, **** P≤0.0001).

### Reduced *ex vivo* insulin-stimulated glucose uptake in fetal skeletal muscle exposed to maternal WSD is associated with attenuated insulin signaling

IRS1 and p110α abundance were significantly decreased in the maternal WSD group (Figure 3A-C) while insulin receptor, p85α, Akt, GSK3β, and GLUT1 abundance were not different between groups (Supplementary Figure 1). While insulin-stimulated IRS1(Y896) phosphorylation in fetal muscle was not affected by maternal diet, Akt phosphorylation at T308, and S473 was significantly lower in WSD compared to CON (Figure 3D-F, I). Consistent with reduced insulin-stimulated Akt activation, AS160, and GSK3β phosphorylation showed no insulin-stimulated increase over basal in the WSD group (Figure 3G-I). These data suggest that developmental exposure to maternal obesity and a WSD reduces insulin action in fetal muscle *ex vivo* even in the absence of systemic factors.

**Figure 3:**
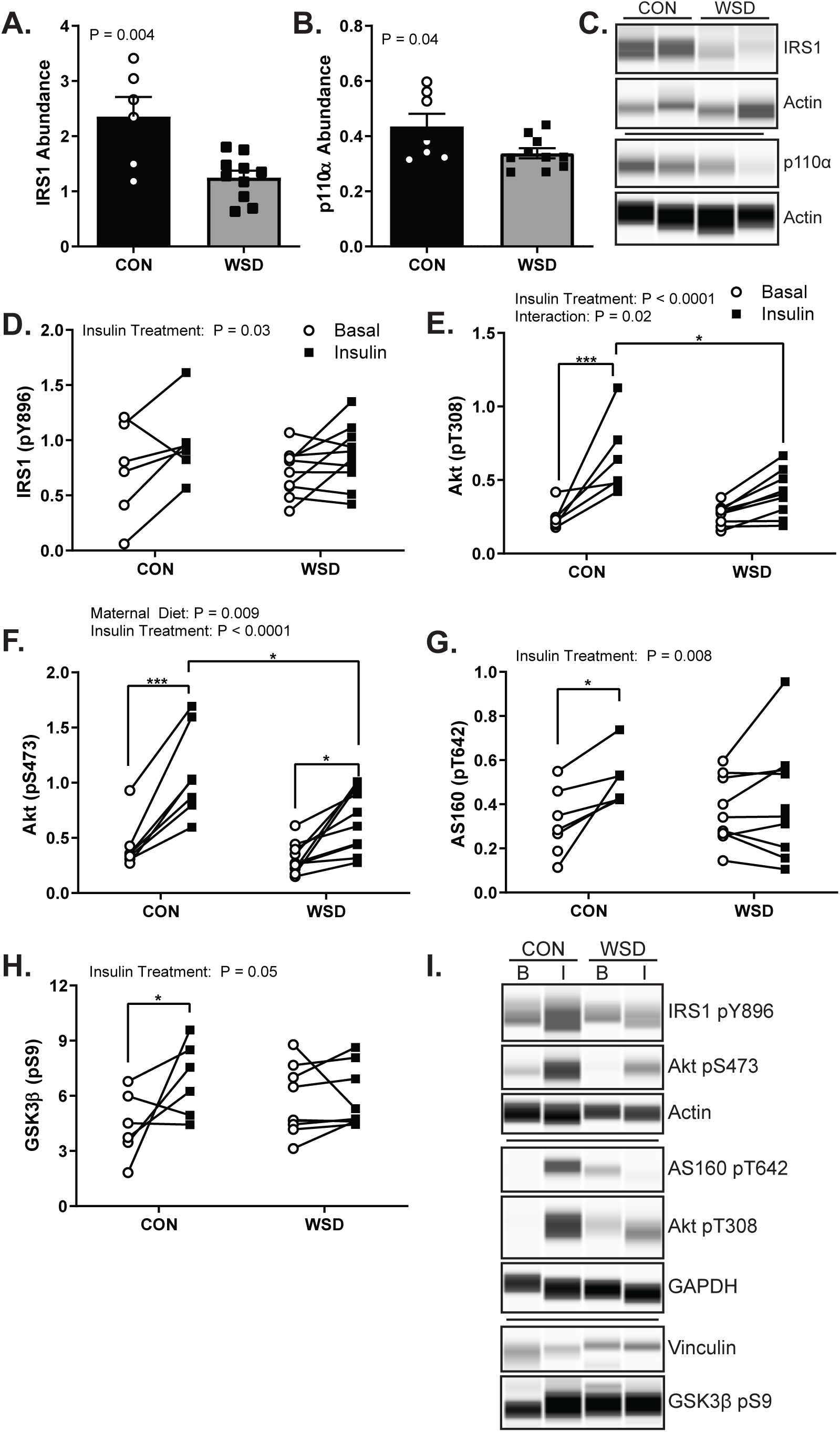
Fetal Skeletal Muscle *Ex Vivo* Insulin Response. The abundance of total IRS1 (A) and total p110α (B) measured in basal soleus muscle homogenate expressed as the peak area relative to the control protein Actin. Example of simple western probing for IRS1, p110α, and Actin (C). Quantification of simple western probing of soleus muscle homogenates for insulin responsive phosphorylation of IRS1 Y896 (D), Akt T308 (E), Akt S473 (F), AS160 T642 (G), and GSK3β S9 (H) expressed as the peak area relative to control proteins Actin, GAPDH, or Vinculin in basal (○) and insulin-stimulated (▪) samples *ex vivo*. Example simple western probing for insulin responsive protein phosphorylation (I). Data is expressed as mean ± SEM, with individual data points shown. Total protein abundance was analyzed by unpaired T test. Phospho-Akt abundance was analyzed by two-way ANOVA with Tukey multiple comparisons test. All other phospho-protein abundance was analyzed by mixed effect model with Sidak correction.

### Juvenile phenotype

A separate cohort of offspring were studied at approximately 7 months of age, prior to weaning, and again at 14 months of age. At ∼8 months of age, offspring were weaned to new group housing and either maintained on the same diet as their maternal group or switched to the opposite diet creating four diet groups (maternal diet/offspring diet: CON/CON, CON/WSD, WSD/CON and WSD/WSD; Figure 1). At 13 months of age, offspring exposed to maternal WSD had increased body weight relative to the offspring from the maternal CON group (Table 2). Post-weaning WSD also resulted in increased body weight, and decreased glucose AUC. Neither maternal nor post-weaning WSD resulted in increases in body fat percentage, fasting insulin, fasting glucose, or IAUC in juvenile offspring. The phenotype of a larger cohort of 14 month old juvenile offspring has been previously reported (15,26). In the larger cohort, fasting insulin and insulin AUC were increased in the WSD/WSD group. In our subset, fasting insulin trended towards being increased in the WSD/WSD animals, but did not reach significance.

**Table 2:**
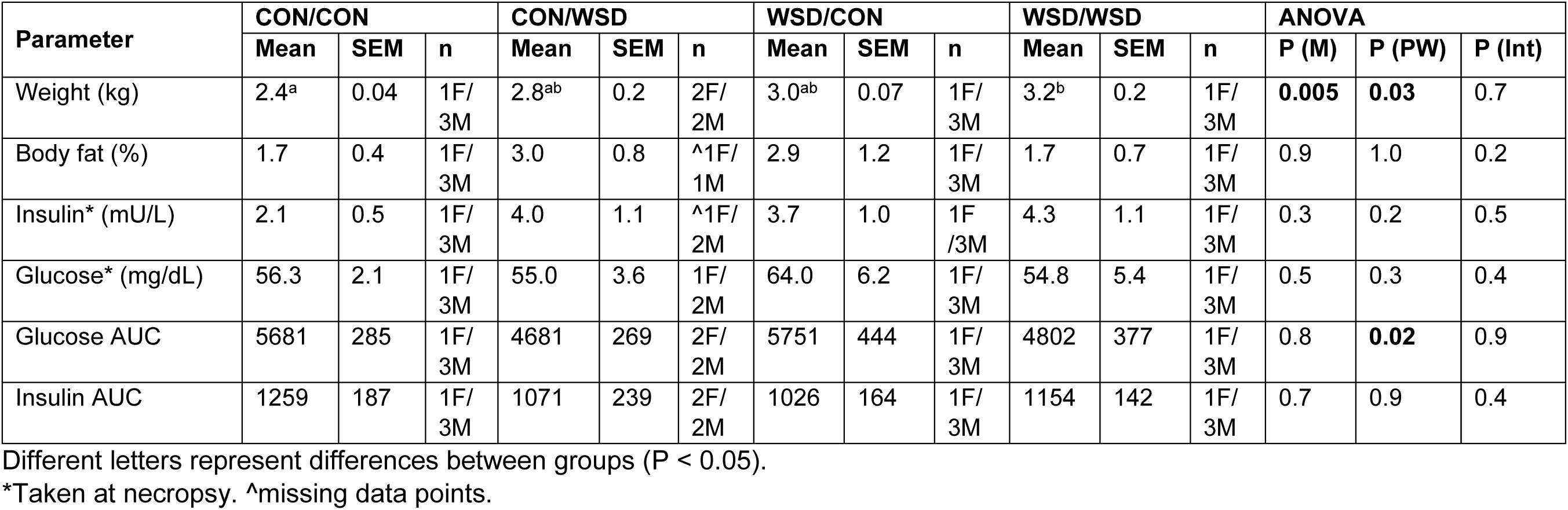
Juvenile Phenotype at age 14 mo of age.

| Parameter | CON/CON |  |  | CON/WSD |  |  | WSD/CON |  |  | WSD/WSD |  |  | ANOVA |  |  |
| --- | --- | --- | --- | --- | --- | --- | --- | --- | --- | --- | --- | --- | --- | --- | --- |
|  | Mean | SEM | n | Mean | SEM | n | Mean | SEM | n | Mean | SEM | n | P (M) | P (PW) | P (Int) |
| Weight (kg) | 2.4<sup>a</sup> | 0.04 | 1F/<br>3M | 2.8<sup>ab</sup> | 0.2 | 2F/<br>2M | 3.0<sup>ab</sup> | 0.07 | 1F/<br>3M | 3.2<sup>b</sup> | 0.2 | 1F/<br>3M | **0.005** | **0.03** | 0.7 |
| Body fat (%) | 1.7 | 0.4 | 1F/<br>3M | 3.0 | 0.8 | <sup>^</sup>1F/<br>1M | 2.9 | 1.2 | 1F/<br>3M | 1.7 | 0.7 | 1F/<br>3M | 0.9 | 1.0 | 0.2 |
| Insulin* (mU/L) | 2.1 | 0.5 | 1F/<br>3M | 4.0 | 1.1 | <sup>^</sup>1F/<br>2M | 3.7 | 1.0 | 1F/<br>3M | 4.3 | 1.1 | 1F/<br>3M | 0.3 | 0.2 | 0.5 |
| Glucose* (mg/dL) | 56.3 | 2.1 | 1F/<br>3M | 55.0 | 3.6 | 1F/<br>2M | 64.0 | 6.2 | 1F/<br>3M | 54.8 | 5.4 | 1F/<br>3M | 0.5 | 0.3 | 0.4 |
| Glucose AUC | 5681 | 285 | 1F/<br>3M | 4681 | 269 | 2F/<br>2M | 5751 | 444 | 1F/<br>3M | 4802 | 377 | 1F/<br>3M | 0.8 | **0.02** | 0.9 |
| Insulin AUC | 1259 | 187 | 1F/<br>3M | 1071 | 239 | 2F/<br>2M | 1026 | 164 | 1F/<br>3M | 1154 | 142 | 1F/<br>3M | 0.7 | 0.9 | 0.4 |

Different letters represent differences between groups (P < 0.05).
\*Taken at necropsy. <sup>^</sup>missing data points.

### Reduced insulin-stimulated skeletal muscle glucose uptake in offspring exposed to maternal WSD persists in 1-year old juveniles

To determine if decreases in insulin-stimulated glucose uptake in fetal skeletal muscle of WSD fed mothers persists and influences the effects of post-weaning WSD, we measured 2DG uptake in gastrocnemius and soleus muscles of 14 month old offspring. In the gastrocnemii, at both insulin concentrations, post-weaning WSD reduced 2DG uptake within the maternal CON groups (Figure 4A, B). Within the maternal WSD groups, post-weaning WSD reduced 2DG uptake as compared to CON/CON (maternal CON/post-weaning CON) but was not different from offspring exposed only to maternal WSD or only to post-weaning WSD. In the soleus muscle, maternal WSD resulted in reduced insulin-stimulated 2DG uptake within the post-weaning CON groups at both insulin concentrations (Figure 4C, D). There was no difference between post-weaning diet within each maternal group at the lower insulin dose. At the higher insulin concentration, post-weaning WSD led to significantly reduced insulin-stimulated 2DG uptake within the maternal CON group. Again, there was no difference between post-weaning WSD and CON in offspring exposed to maternal WSD. Taken together, this data shows that maternal WSD exposure results in defects in insulin-stimulated 2DG uptake that persists in skeletal muscle post-weaning at one year of age prior to increases in offspring adiposity or clinical measures of impaired glucose tolerance. The magnitude of suppression is similar to that of a post-weaning WSD alone.

**Figure 4:**
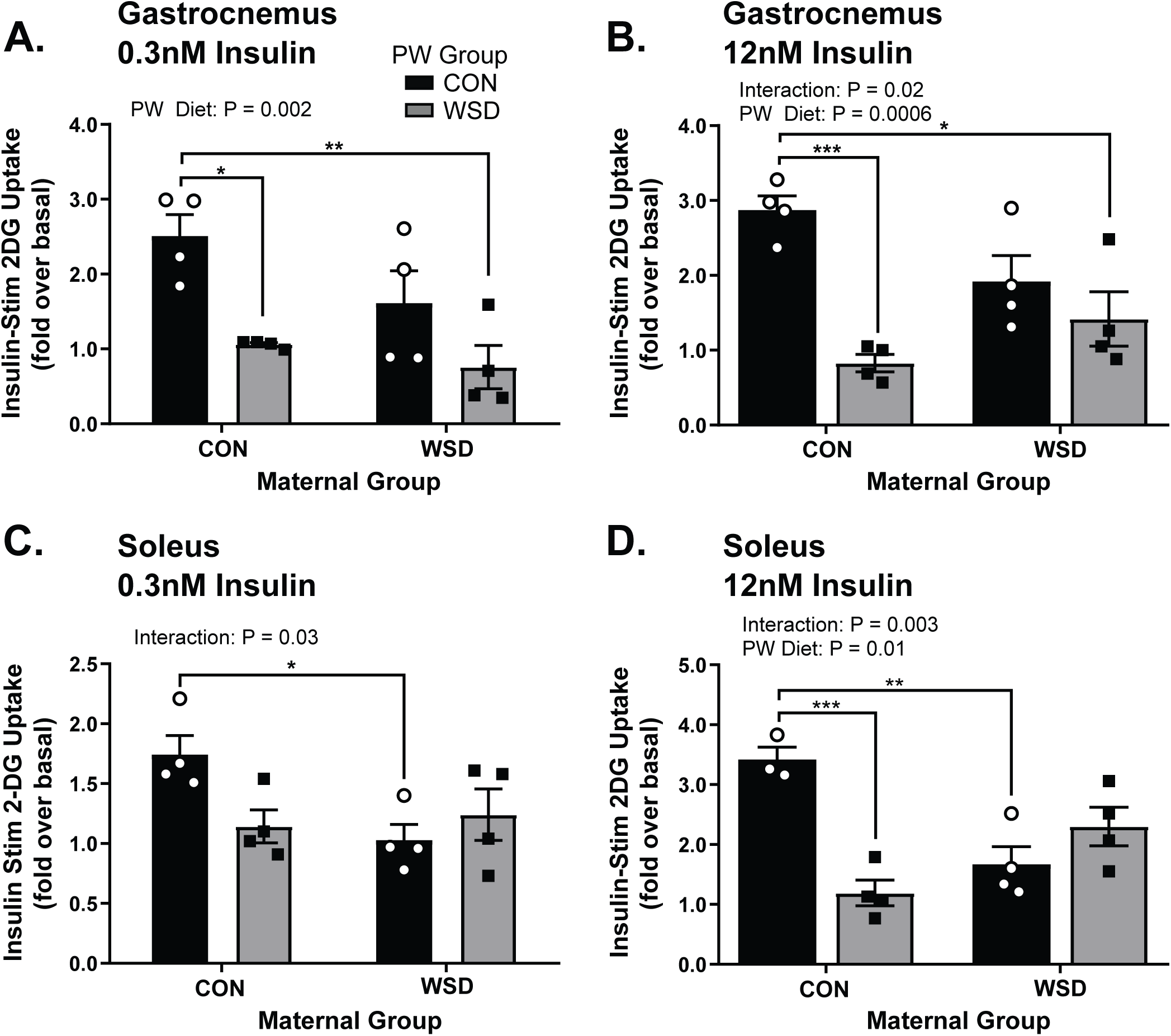
Juvenile Skeletal Muscle *Ex Vivo* Glucose Uptake. Insulin-stimulated 2-DG uptake measured *ex vivo* in 1-year old juvenile gastrocnemius at 0.3nM (A) or 12nM (B) insulin concentration. Insulin-stimulated 2-DG uptake measured in the soleus muscle of 1-year old NHP *ex vivo* at 0.3nM (C) or 12nM (D) insulin concentration. Insulin-stimulated 2-DG uptake is reported as a fold increase over basal (unstimulated) glucose uptake. Data is expressed as mean ± SEM, with individual data points shown, and analyzed by two-way ANOVA with Tukey multiple comparisons test. Brackets indicate group comparisons (* P≤0.05, ** P≤0.01, *** P≤0.001).

### Activation of insulin signaling is impaired in skeletal muscle of juvenile offspring exposed to maternal WSD

To determine the mechanisms for reduced insulin-stimulated 2DG uptake, the abundance and activation state of key insulin signaling intermediates was measured in basal and insulin-stimulated (12nM) gastrocnemius. There was no difference in the total abundance of IRS1 or p110α as previously observed in the fetal muscle or in Akt by maternal or post-weaning diet (Supplementary Figure 2). We observed a significant effect of diet group on IRS1 phosphorylation by two-way ANOVA, however, there was not a specific difference in insulin effect among groups (Figure 5A, E). There was, however, a reduction in insulin-stimulated phosphorylation of Akt at S473, but not T308, in all groups relative to CON/CON (Figure 5B, C, E). The lack of a significant difference between basal and insulin-stimulated Akt pS473 in all diet groups other than CON/CON corresponds to observed reductions in insulin stimulation of glucose uptake. Insulin stimulation of AS160 phosphorylation at T642 also showed a significant effect by insulin treatment, but not by group (Figure 5D, E). Together, we find that insulin signaling activation was reduced by either maternal or post-weaning WSD.

**Figure 5:**
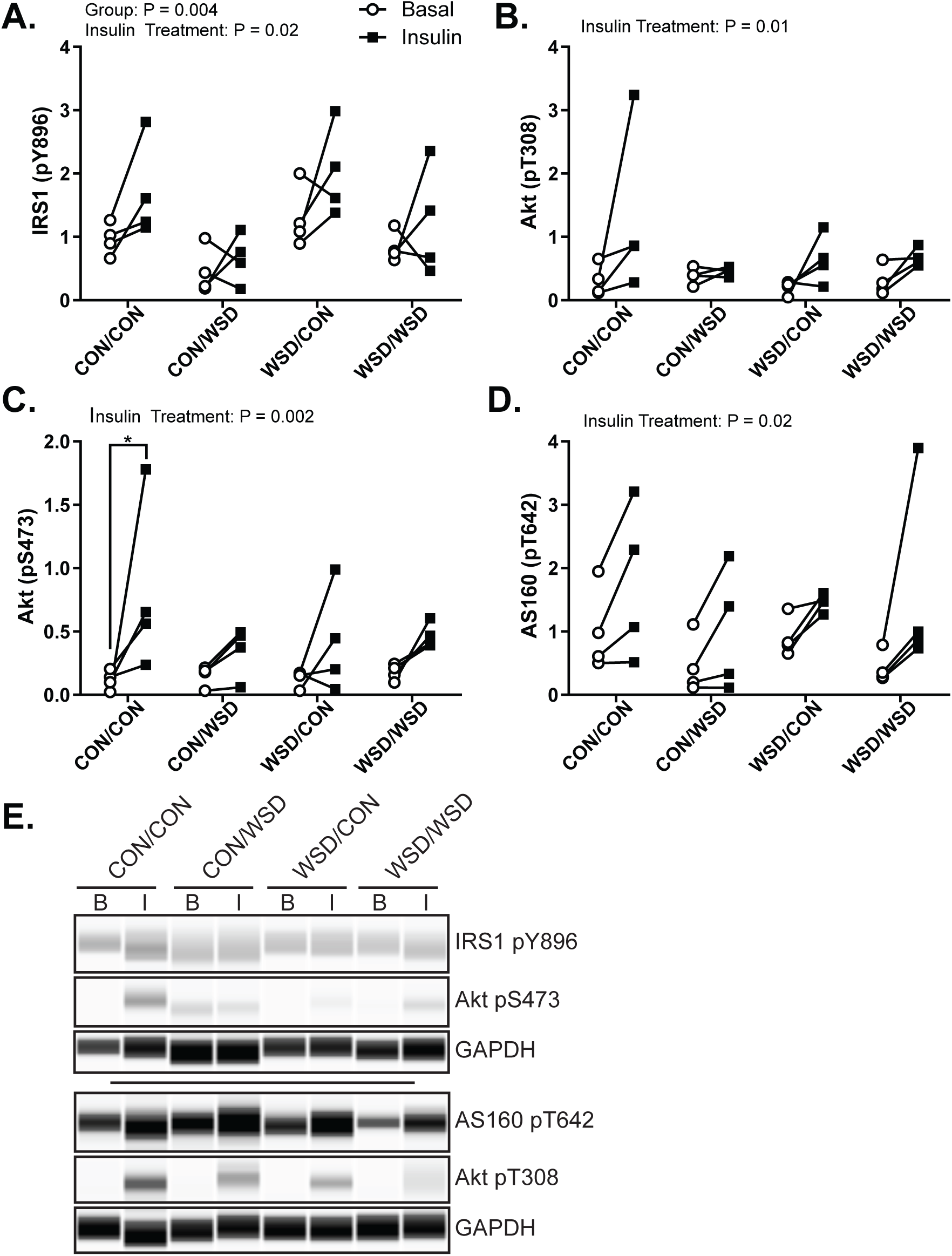
Juvenile Skeletal Muscle *Ex Vivo* Insulin Response. Insulin responsive protein phosphorylation measured by simple western in *ex vivo* basal (○) and insulin-stimulated (▪) soleus. IRS1 pY896 (A), Akt pT308 (B), Akt pS473 (C), AS160 pT642 (D) peak area expressed as a ratio to the control protein GAPDH. Example simple western of phospho-protein staining in basal (B) or insulin-stimulated (I) samples (E). CON/CON – Maternal CON diet, post-weaning CON diet group; CON/WSD – Maternal control diet, post-weaning WSD group; WSD/CON – Maternal WSD, post-weaning CON diet group; WSD/WSD – Maternal WSD, post-weaning WSD group. Data was analyzed by two-way ANOVA with Tukey multiple comparisons test. Brackets indicate group comparisons (* P≤0.05).

### Activation of the skeletal muscle insulin signaling cascade in vivo in offspring is reduced in association with exposure to a maternal WSD

To complement our *ex vivo* measures of skeletal muscle insulin signaling, we also measured activation of the insulin signaling cascade in soleus muscle biopsies from 7 month old animals before and after an intravenous insulin bolus. For reference, offspring start eating independently by 4 months of age (27); therefore, data collected in 7 month offspring reflects a combination of maternal exposures during fetal development through lactation and WSD exposure through food introduction during this pre-weaning developmental period. In offspring muscle, phosphorylation of both IRS1 and Akt was significantly reduced in insulin-stimulated muscle of animals from the maternal WSD group (Figure 6A, B, F). Insulin-stimulated GSK3β phosphorylation was increased in the WSD but not CON (Figure 6C, F). The increase in GSK3β activation corresponded to a significant increase in the total abundance of GSK3β after insulin stimulation indicating that the difference in activation between CON and WSD is not due solely to an increase in the fraction of GSK3β that is phosphorylated (Figure 6D, F). Despite differences in Akt phosphorylation, insulin treatment resulted in a significant effect on AS160 phosphorylation that was not different by maternal diet group (Figure 6E, F). Consistent with the ex vivo juvenile signaling data, but in contrast to the fetal insulin signaling data, we observed no difference in total abundance of IR, IRS1, p110α, or Akt (Supplementary Figure 3).

**Figure 6:**
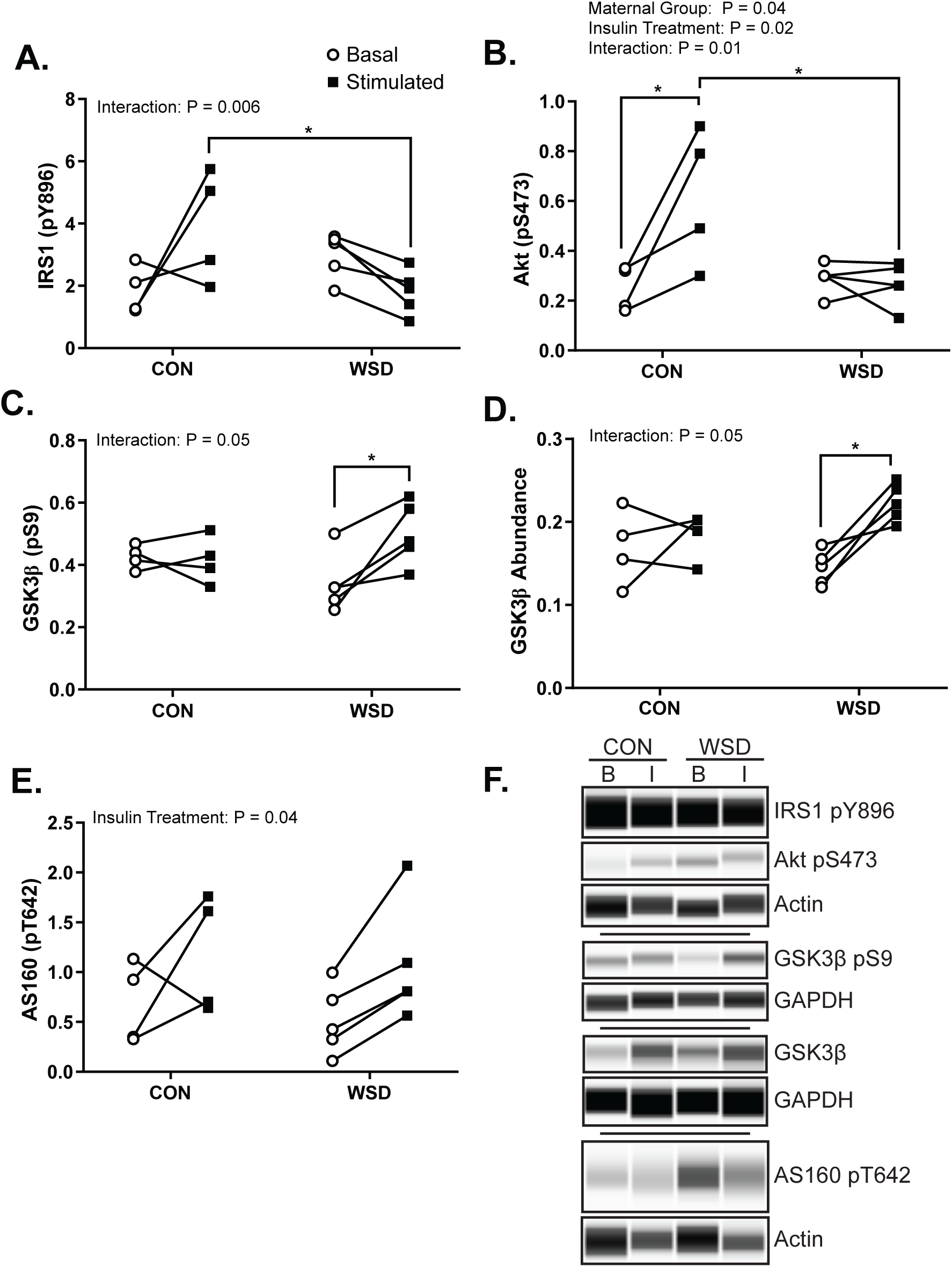
Juvenile Skeletal Muscle In Vivo Insulin Response. Insulin responsive protein phosphorylation measured by simple western in gastrocnemius biopsies collected before (Basal ○) and 10min after (Stimulated ▪) intravenous insulin injection. IRS1 pY896 (A), Akt pS473 (B), GSK3β pS9 (C), total GSK3β (D), AS160 pT642 (E). Abundance expressed as a ratio of protein peak area relative to Actin or GAPDH. Example simple western probing of protein abundance (F). Data was analyzed by two-way ANOVA with Tukey multiple comparisons test. Brackets indicate group comparisons (* P≤0.05).

## Discussion

Maternal obesity is strongly associated with greater infant adiposity, insulin resistance (5), and an earlier risk for developing metabolic diseases like type 2 diabetes and nonalcoholic fatty liver disease (28–30); however, few studies have directly examined the effect of maternal obesity and WSD on activation of insulin signaling pathways in offspring tissues regulating glucose uptake and utilization, such as skeletal muscle. Skeletal muscle insulin resistance and impaired glucose uptake is a common metabolic disorder in obese individuals and a primary contributor to the etiology of type 2 diabetes (11,12). Here, we investigated insulin stimulation of glucose uptake in skeletal muscle of offspring exposed to maternal obesity and WSD during early developmental periods (i.e. fetal and pre-weaning) and shortly after weaning in Japanese macaques. We show for the first time that intrauterine exposure to maternal WSD induced obesity reduces insulin-stimulated glucose uptake and activation of insulin signaling in fetal skeletal muscle in association with a reduction in IRS-1 and p110α.

Fetal exposure to maternal WSD reduced fetal skeletal muscle response to insulin stimulation of glucose uptake at both low and high insulin concentrations in soleus and to a lesser extent in the rectus femoris. Interestingly, basal glucose uptake was increased in the soleus. Reduced glucose uptake in fetal muscle was accompanied by attenuated expression of IRS1 and the p110α subunit of PI3K, potentially resulting in reduced capacity to respond to insulin, or reduced sensitivity. Consistent with reduced insulin signaling activation, insulin-stimulated phosphorylation of Akt and substrates of Akt were reduced by maternal obesity and WSD. Surprisingly, we saw no difference by maternal diet in insulin-stimulated phosphorylation of IRS1 in the fetal muscle. Impaired signaling at the level of IRS1 activation is a common marker of insulin resistance with high fat diet or obesity in adults and is often linked to elevated activation of pro-inflammatory cytokine or stress signaling by JNK or atypical PKCs (31). Our observations in fetal samples suggests that glucose transport is attenuated by maternal obesity through reduced signal transduction prior to Akt activation, potentially through reduced PI3 Kinase activity.

In agreement with our fetal data, maternal obesogenic diet during pregnancy and lactation resulted in reduced abundance of both IRS1 and the p110β subunit of PI3K in adult rodent offspring skeletal muscle (32) and adipose tissue (33). Further, IRS1-PI3K activity, but not IRS1 phosphorylation, was reduced in fetal muscle in offspring of obese sheep (34); although, these measures in fetal sheep were made in fasted, and not insulin-stimulated tissues, it provides additional evidence that IRS1-PI3K may be a critical node regulated by maternal obesity in the fetal muscle. Lastly, reduced insulin-stimulated glucose uptake in fetal muscle has also been found in a rat model of fetal hyperglycemia with impairments in insulin signaling occurring at the level of Akt and not IRS1 (35).

The decrease in glucose uptake may also be associated with low oxidative capacity and glucose metabolism reported previously in primary myotubes from WSD fetuses. Previously we found that fetal muscle responds to maternal obesity and WSD by upregulating fatty acid metabolism, mitochondrial complex activity, and metabolic switches (CPT-1, PDK4) that promote lipid utilization over glucose oxidation (17). The mechanisms underpinning this adaption are unclear but are likely initiated by functional demands on the muscle given the primary energy-rich high fat diet and reduced oxygen supply (36). Several studies have also demonstrated a relationship between mitochondrial dysfunction and muscle cell insulin function at the level of IRS1 abundance, therefore, the previously observed changes in mitochondrial function induced by maternal WSD may also contribute to loss of insulin response observed here (37,38).

We also observed persistent insulin resistance in skeletal muscle glucose transport of otherwise healthy 14 month old juvenile offspring from obese WSD dams despite weaning offspring to a healthy control diet. Insulin-stimulated glucose uptake was reduced in the soleus, and to a lesser extent in the gastrocnemius, of offspring exposed to maternal WSD but weaned to a healthy diet. All three groups exposed to WSD exhibited a loss of insulin-stimulated Akt phosphorylation relative to the CON/CON group. Given the modest signaling response observed *ex vivo*, we also assessed the activation of the insulin signaling cascade in skeletal muscle biopsies from 7 month old offspring before and after intravenous administration of an insulin bolus. Offspring of WSD dams had reduced insulin activation of both IRS1 and Akt in agreement with our *ex vivo* signaling data. Downregulation in IRS1 activation may reflect exposure to postnatal WSD through a combination of independent eating and lactation. Unexpectedly, activation of GSK3β and AS160, downstream targets of Akt, were unaffected by maternal WSD. We speculate that this may be due to a compensatory increase in protein abundance in response to insulin as seen with GSK3β in WSD offspring. Together, these observations suggest that the mechanism of insulin resistance in the fetal skeletal muscle occurs downstream of IRS1 and is different from obesity or WSD-induced insulin resistance in juvenile animals.

Although increasing evidence indicates that developmental exposures to maternal obesity, diabetes or a poor-quality maternal diet increases offspring risk of obesity and metabolic diseases, the mechanisms that drive this phenomenon are poorly understood. Many of the metabolic adaptations in offspring are thought to be driven by changes in placental lipid transport (21,39,40), reduced oxygen delivery to the developing fetus (36,41) or increased fetal exposure to inflammatory cytokines (42). Fetal adaptations to this altered in utero environment are thought to persist into the postnatal period due, in part, to changes in the offspring epigenome, the activity of epigenetic regulators, which can influence the expression of molecules important to hormone signaling and metabolism, and/or through changes in noncoding RNAs such as microRNAs and long noncoding RNAs (43–45).

While our study is unique in examining functional measures of insulin activation in skeletal muscle, our non-human primate model limits us to observational data. In the absence of loss- and gain-of function studies, we cannot conclude that the molecular changes detected are causative of reduced insulin-stimulated glucose uptake. Additionally, in this small cohort, we cannot separate out the effects of maternal WSD from that of maternal obesity or detect differences between male and female offspring. Despite these limitations, our conclusions are based on direct measures of insulin-stimulated glucose transport and signaling pathway activation. While this study has focused on insulin activation of skeletal muscle in response to maternal obesity and WSD, it should be noted that all of the changes found in the juvenile offspring occurred in the absence of overt adiposity or hyperinsulinemia. Thus, the reduced skeletal muscle glucose transport and phosphorylation preceded whole-body insulin resistance. Evidence for fetal programming of adult body composition and insulin resistance in humans is derived mostly from epidemiological studies, associating poor prenatal nutrition/obesity with increased fat mass, central distribution of fat, and metabolic syndrome (46).

Previous work in this model has shown that maternal obesity and WSD results in greater hepatic lipid accumulation and de novo lipid synthesis in fetal and postnatal offspring at 14mo (16,17,36) and significantly elevated β:α cell ratio in offspring at 14 and 36mo of age (15,47). Importantly, in viewing our outcomes in the context of changes within the physiologic system, it is easy to predict that these offspring would be more susceptible to earlier development of whole body insulin resistance and metabolic diseases when faced with challenges like a continued WSD, sedentary lifestyle or even puberty.

Together, these data are the first to suggest that maternal obesity with WSD exposure *in utero* results in reduced skeletal muscle insulin response that persists at one year of age and is comparable to post-weaning WSD exposure alone. Given that the offspring were young and lean with normal glucose tolerance, we speculate that these are primary changes, involved in the development of muscle insulin resistance that may manifest themselves during puberty or future metabolic challenges. While we have observed changes in insulin response and insulin-stimulated glucose uptake driven by maternal obesity and WSD, the mechanism for this altered response still remains to be determined and is an important goal of our future studies.

## Author Contributions

C.E.M. is responsible for the conception and design of the experiments. C.E.M., B.H., W.C.B., S.S., S.R.W., D.L.T., and T.A.D. collected the samples, conducted the research and analyzed data. J.E.F., K.M.A., E.L.S., P.K., D.L.T., and T.A.D. maintained and supported the animal model. B.H., W.C.B. and C.E.M. wrote the manuscript. S.S., S.R.W., K.M.A., E.L.S., P.K., M.G. and J.E.F. contributed to the interpretation of the data and revision of the manuscript. All authors read and approved the final manuscript. C.E.M is the guarantor for this work and, as such, had full access to all the data in the study and takes responsibility for the integrity of the data and the accuracy of the data analysis.

## Prior Presentation

Parts of this study were presented in abstract format at the 76th Scientific Sessions of the American Diabetes Association, June 10-14, 2016, New Orleans.

