## Supplemental Figure 1 for "Maternal Obesity and Western-style Diet Impair Fetal and Juvenile Offspring Skeletal Muscle Insulin-Stimulated Glucose Transport in Nonhuman Primates"

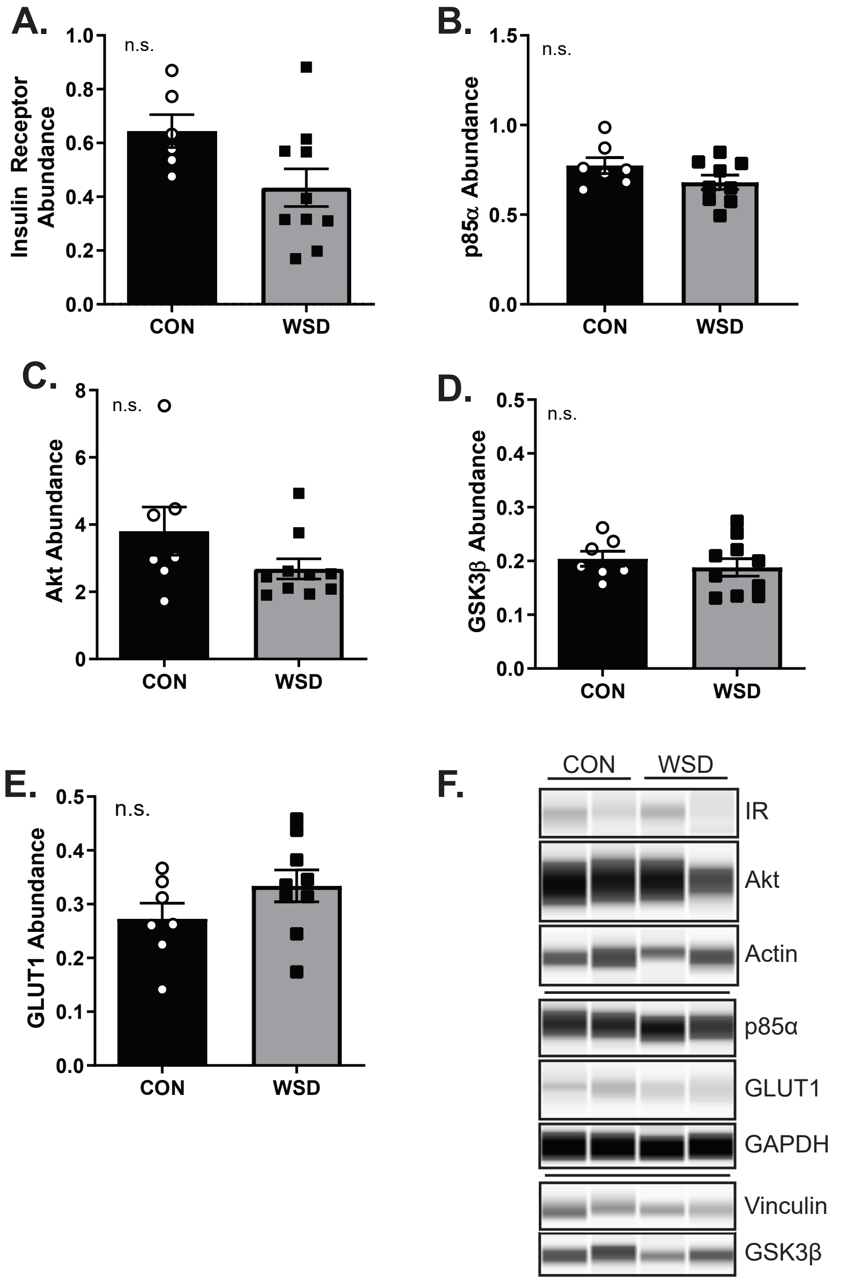


**Supplementary Figure 1:** Fetal Skeletal Muscle Insulin Signaling. Total abundance of components of the insulin signaling pathway measured by simple western in homogenates of fetal soleus muscle. (A) Insulin receptor abundance as a ratio to the control protein Actin. (B) p85α abundance as a ratio to the control protein GAPDH. (C) Akt abundance relative to the control protein Actin. (D) GSK3β abundance relative to the control protein Vinculin. (E) GLUT1 abundance relative to the control protein GAPDH. (F) Example simple western probing of each of the above proteins. Data is expressed as mean ± SEM, with individual data points shown, and analyzed by unpaired T test.
