## Supplemental Figure 2 for "Maternal Obesity and Western-style Diet Impair Fetal and Juvenile Offspring Skeletal Muscle Insulin-Stimulated Glucose Transport in Nonhuman Primates"

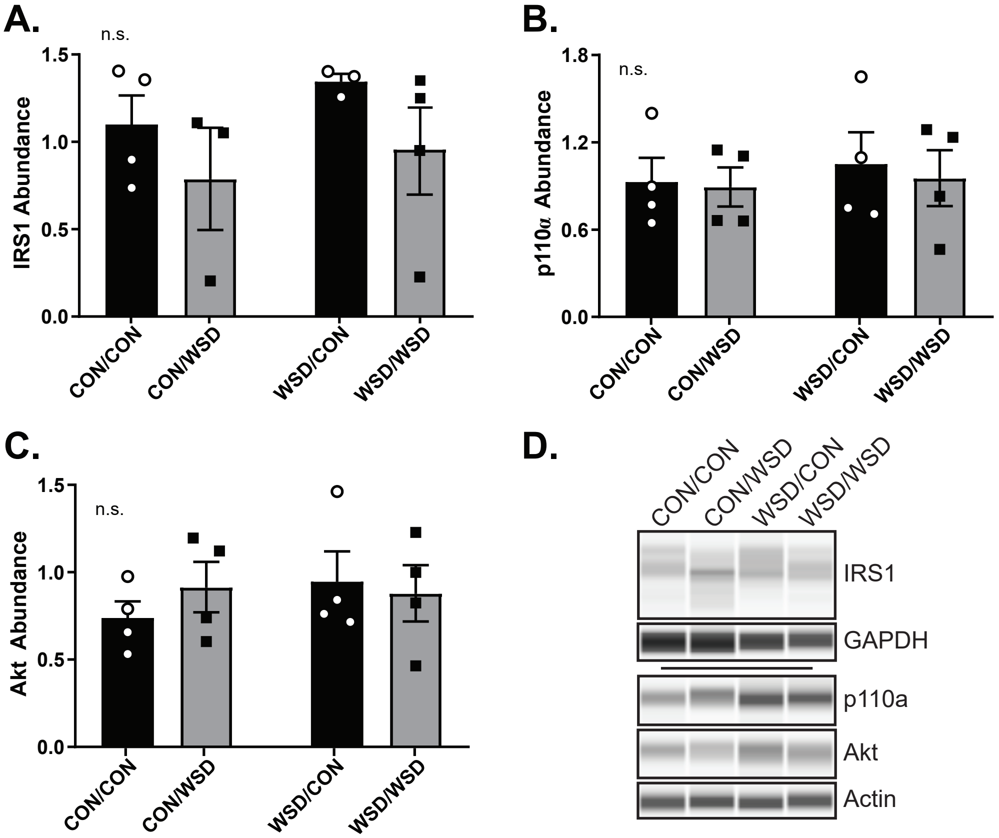


**Supplementary Figure 2:** Juvenile Skeletal Muscle Ex Vivo Insulin Response. Total abundance of proteins in the insulin signaling cascade were measured in basal skeletal muscle homogenates measured by simple western. (A) IRS1 abundance is expressed relative to GAPDH. (B) p110α abundance is expressed relative to Actin. (C) Akt abundance is expressed relative to Actin. (D) Example simple western probing of IRS1, p110α, Akt, GAPDH, and Actin. Data is expressed as the mean ± SEM, with individual data points shown, and analyzed by two-way ANOVA with Tukey multiple comparisons test.
