## Supplemental Figure 3 for "Maternal Obesity and Western-style Diet Impair Fetal and Juvenile Offspring Skeletal Muscle Insulin-Stimulated Glucose Transport in Nonhuman Primates"

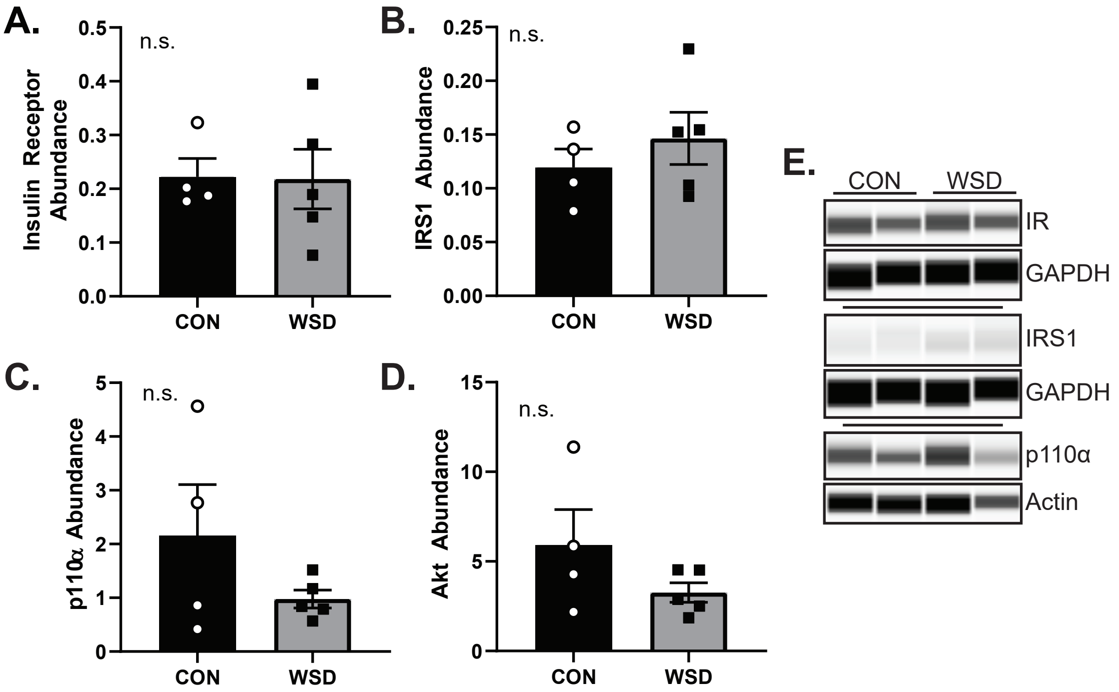


**Supplementary Figure 3:** Juvenile Skeletal Muscle In Vivo Insulin Response. Abundance of total proteins in the insulin signaling cascade were measured by simple western in homogenates of gastrocnemius biopsies from fasted seven-month-old animals. Insulin receptor (A), IRS1 (B), p110α (C), and Akt (D) abundance was expressed relative to either GAPDH or Actin. Data is expressed as the mean ± SEM, with individual data points shown, and analyzed by unpaired T test.
