## Supplemental Table 1 for "Maternal Obesity and Western-style Diet Impair Fetal and Juvenile Offspring Skeletal Muscle Insulin-Stimulated Glucose Transport in Nonhuman Primates"

### Supplementary Table 1

| Target | Manufacturer | Catalog Number | Dilution |
|---|---|---|---|
| IRS1 | Cell Signaling Technology | 3407 | 1:25 |
| p110α | Cell Signaling Technology | 4249 | 1:50 |
| IRS1 (pY896) | Abcam | ab46800 | 1:25 |
| Akt (pT308) | Cell Signaling Technology | 4056 | 1:50 |
| Akt (pS473) | Cell Signaling Technology | 4060 | 1:50 |
| AS160 (pT642) | Cell Signaling Technology | 4288 | 1:25 |
| GSK3β (pS9) | Cell Signaling Technology | 9323 | 1:100 |
| Insulin Receptor | Cell Signaling Technology | 3025 | 1:20 |
| p85α | Cell Signaling Technology | 4257 | 1:25 |
| Akt | Cell Signaling Technology | 4691 | 1:1000 |
| GSK3β | Cell Signaling Technology | 9315 | 1:2000 |
| GLUT1 | Cell Signaling Technology | 12939 | 1:25 |
